# The Role of Gene Encoding Variation of *DRD4* in the Relationship between Inattention and Seasonal Daylight

**DOI:** 10.1101/825083

**Authors:** M.A. Vollebregt, B. Franke, J.K. Buitelaar, L.E. Arnold, S.V. Faraone, E.H. Grevet, A. Reif, T. Zayats, J. Bralten, C.H.D. Bau, J. Haavik, J. Kuntsi, R.B. Cupertino, S.K. Loo, A.J. Lundervold, M. Ribasés, C. Sánchez-Mora, J.A. Ramos-Quiroga, P. Asherson, J.M. Swanson, M. Arns

## Abstract

Daylight is the strongest synchronizer of human circadian rhythms. The circadian pathway hypothesis posits that synchrony between daylight and the circadian system relates to (in)attention. The dopamine neurotransmitter system is implicated in regulating the circadian system as well as in (attention)-deficit hyperactivity disorder [ADHD]. We studied the role of functional genetic variation in the gene encoding of dopamine-receptor-D4 (*DRD4*) in the relationship between inattention and seasonal daylight (changes). Gene-by-environment (GxE) mega-analyses were performed across eight studies including 3757 adult participants (with and without ADHD). We tested 1) the *Spring-focus hypothesis*, in which attention in 7R-carriers normalizes with increasing daylight levels preceding measurement, 2) the *Summer-born ADHD hypothesis*, in which 7R-carriers report more inattention when born in spring/summer than in autumn/winter, 3) the *Winter-born ADHD hypothesis*, opposing the second hypothesis. The *Spring-focus hypothesis* was upheld (1386 ADHD, 760 controls; *d*=-0.16 between periods); 7R-carriers reported even *less* inattention than 7R-non-carriers after winter solstice (*d*=0.27 between genotype-groups). Results were diagnosis-independent. Sensitivity analyses at individual study level confirmed the circannual patterns for 7R-carriers. Incorporating geographic changes into the independent measure, we also calculated changes in sunlight levels. This approach likewise showed that inattention correlated negatively with increasing light levels in 7R-carriers (*r*=-.135). Results emphasize peripheral effects of dopamine and the effects of (seasonal) daylight changes on cognition.

## INTRODUCTION

### The circadian pathway

Daylight is the strongest synchronizer of human circadian rhythms. When daylight reaches the retina, it provides the internal clock system [suprachiasmatic nuclei (SCN)] with information about the time of day, thereby leading to daylight entrainment (1). Even modest misalignment of the internal clock from sleep/wake behavior can result in poorer sleep quality (2). Several lines of evidence support the idea that sleep deprivation or extended wakefulness is accompanied by a variety in alteration of dopamine signalling (3). Furthermore, shortened sleep duration has been shown to be associated with inattention in healthy individuals (4-6) as well as in people with attention-deficit/hyperactivity disorder (ADHD) (7,8). Multiple studies using various methods have shown that a majority of individuals with ADHD suffer from a circadian phase delay (9-12), with a prevalence as high as 78% in adults (12).

Light-emitting diodes (LED) and the increase in time spent on electronic devices such as computers, smart phones, and tablets compete with daylight as the primary cue that entrains the biological clock to a 24-hour (24h) rhythm. The ‘circadian pathway’ hypothesis posits that artificial blue light exposure in the evening delays sleep onset, thereby reducing sleep duration, which, in turn, results in increased symptoms of inattention (13,14). Each step in this pathway has recently been confirmed in a sample of school-aged children with ADHD (14). The phase-delaying effects of artificial blue light exposure in the evening can be counteracted by intense natural light in the morning (15), when our circadian clock is most sensitive to entrainment to the 24h rhythm (16).

Exposure to intense natural light in the morning is more common in geographic areas characterized by high sunlight intensity. Prevalence rates of ADHD are lower in these areas compared to those with less sunlight intensity (17,18).

### Gene-specific responses within the circadian pathway

Although many environmental risk factors for ADHD have been studied (19), circannual factors such as (changes in) daylight exposure alone and its interaction with genotypic variation have scarcely been examined. Genotypic variation alone has been investigated in relation to ADHD, one of the first discoveries was an association with the 7-repeat (7R) allele of the *DRD4* gene. This gene has a complex polymorphism in a coding region (exon 3) based on the 48-bp tandem repeats (VNTR), with common alleles defined by 4-repeats (4R), 7-repeats (7R), and 2-repeats (2R) in the population. The first studies [a population-based study (20) and a fail-based study (21)] reported an increased frequency of the 7R allele in children with ADHD, which was subsequently replicated in many studies and multiple meta-analyses (22). Arns et al. (18) proposed that the *DRD4* gene may mediate the earlier observed relationship between sunlight intensity exposure and ADHD prevalence rates. The *D4* receptor is thought to be involved in converting light to neuronal signals in the retina (23), and its transcription exhibits a strong circadian pattern in rodents (24). Cells in the retina that respond to light (photosensitive retinal ganglion cells, pRGCs) express the photopigment melanopsin, and primarily project to the internal clock system (SCN). Stimulating the SCN activates the second messenger cyclic adenosine monophosphate (cAMP), which advances or delays the internal clock (25). Both light and dopamine have an influence on cAMP activation and subsequently on melatonin synthesis (26). With light-sensitive cAMP being an important component of the cyclic rhythmicity of the SCN (27), variation in its formation will affect circadian clock functioning. Asghari et al. (28) showed that rodent ovary cells overexpressing the *DRD4* 7R allele displayed a fluctuation in cAMP that was about twofold reduced compared to other, more common *DRD4* alleles (i.e., 2R and 4R). A study in nearly 700 humans also provided some indication that 7R-carriers reported higher daytime sleepiness than non-carriers (29), further suggesting a relationship between *DRD4* and circadian clock functioning.

The current study focused on genotypic variation in the *DRD4* gene. Based on the work of Asghari et al (28), we initially hypothesized that individuals carrying the *DRD4* 7R allele (7R-carriers) may be less sensitive to light. However, VanderLeest et al. (30) showed that the circadian clock response to light was *enhanced* in (non-genotyped) rodents in controlled light-dark cycles, by exposing them to *short photoperiods* (analogous to human exposure to short winter days) (30-32). In line with these findings, prior light exposure also was shown to alter the way in which the circadian clock aligned to the day/night cycle in humans (33). This led to our *spring-focus hypothesis* (Figure 1a), in which we posit that the circadian clock response to light of 7R-carriers normalizes during a period of increasing light exposure. Studying inattention as an endpoint of the above described ‘circadian pathway’ (13,14), 7R-carriers would be expected to show less inattention in the months following winter solstice, characterized by increasing light exposure. 7R-non-carriers would be expected to be less vulnerable to such subtle seasonal changes in light exposure.

**Figure 1.**
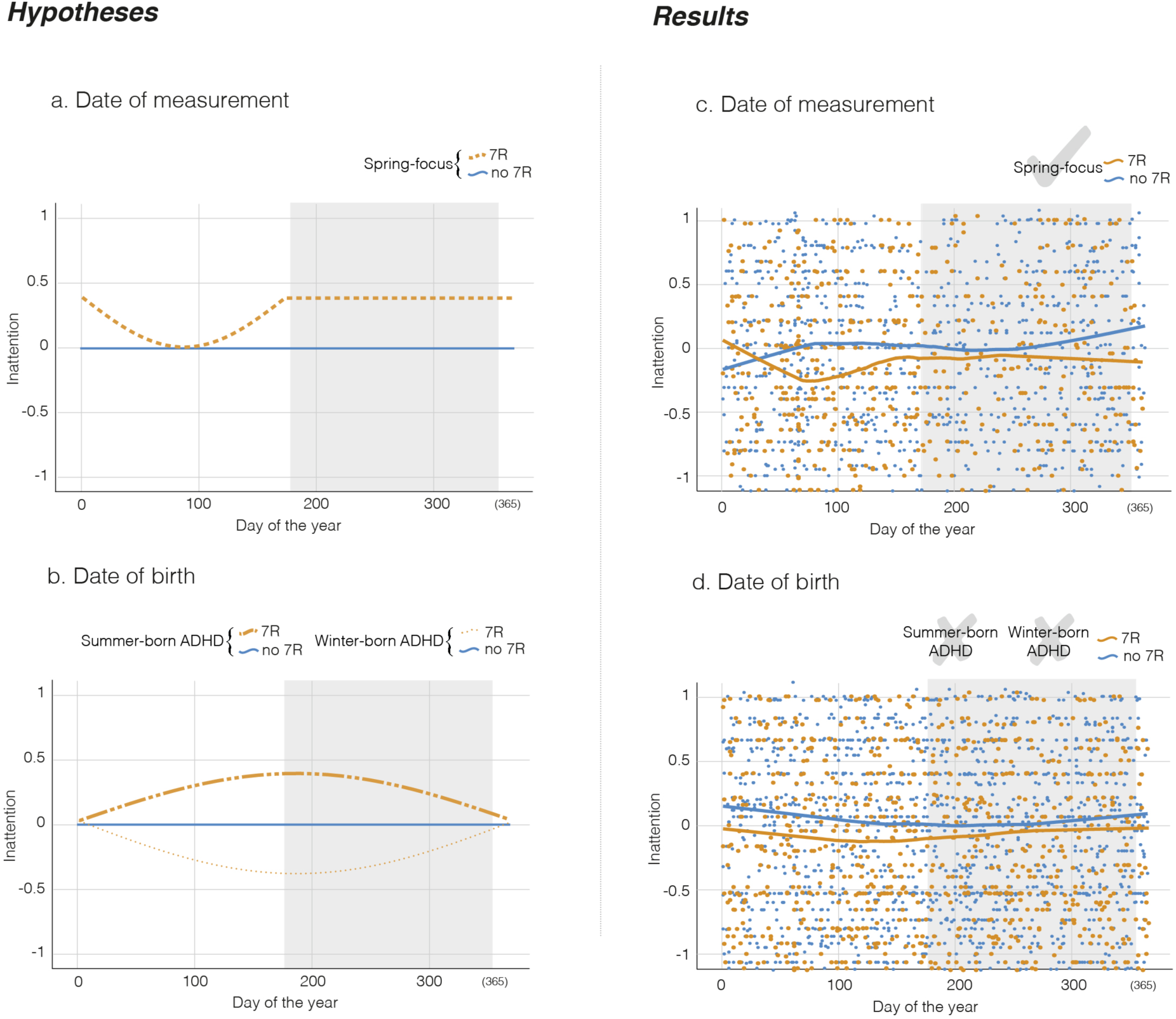
Normalized inattention ratings arranged by a) date of measurement and b) date of birth, according to DRD4 genotype (orange: 7R-carriers, blue: 7R-non-carriers) *(a-b)* as expected by each hypothesis, and *(c-d)* as actually found. The grey-shaded area indicates the period between summer-and winter-solstice, adjusted towards seasonality of the Northern hemisphere, the non-shaded area indicates the analogues period between winter- and summer-solstice. All y-axes are zoomed in to [-1 – 1]. NB: a more negative value implies less inattention, i.e., better attention. The acceptance (checkmark tick) or rejection (cross) of the in *(a-b)* visualized tested hypotheses are displayed in the right top corner of each graph in *(c-d)*. A Loess Fit visualizes variation in the data by nonlinearly comparing data to its neighbouring data. Note that this procedure does not take into account the neighbouring of the last and first day of the year.

### Circadian pathway development

In addition to the acute effects of fluctuation in light exposure, intensity of and seasonal changes after birth have been hypothesized to have phase-sensitive learning effects (also termed ‘imprinting’) on the circadian system (34-36); such phase-sensitive effects have been confirmed for heart rate variability in adulthood (37,38). Likewise, the early lighting environment is thought to shape later adult circadian rhythms (39). After birth, the circadian clock needs to be entrained to exactly 24 hours to prevent misalignment with the external environment, with measurable biological rhythms emerging between 6-18 weeks after birth (40). The effect of seasonal changes in daylight exposure after birth has been studied by comparing chronotypes [morningness (being most active and alert in the morning) and eveningness (being most active and alert in the evening)] in healthy individuals. Eveningness is more likely to occur in individuals born in spring/summer compared to those born in any other season (34,35,41). Yet, another study highlighted more specifically the importance of *changes* in daylight, where eveningness was more common in individuals born in months mostly associated with increasing day length (February to April [August to October in the Southern hemisphere]) compared to months associated with long, decreasing, or short day length (36). Acute effects of sunlight exposure that occur throughout the lifespan may still modulate the enduring effects of phase-sensitive learning after birth. How light environment following birth may shape attentional functioning through circadian system development is yet to be investigated.

### Genotype-specific responses within circadian pathway development

If individuals carrying the *DRD4* 7R allele convert light to neuronal signals in a different way than individuals possessing other *DRD4* alleles, one could hypothesize that the phase-sensitive learning process after birth may be partly genetically determined. Indeed, based on 64 patients and 163 healthy individuals, Seeger and colleagues (42) reported that carrying a *DRD4 7R* allele and being born in spring/summer resulted in a 2.8-fold higher likelihood of children being diagnosed with ‘hyperkinetic disorder’ (overlapping with ADHD) compared with 7R-non-carriers. This led to the *summer-born ADHD hypothesis*, which posits that 7R-carriers born in spring/summer have higher inattention ratings than those born in autumn/winter, while this difference would not be observed in 7R-non-carriers (Figure 1b). Data presented by Brookes et al. (43) are, however, not consistent with the findings of Seeger and co-workers (42). In a sample of 1110 child-parents trios, Brookes et al. found a numerically higher prevalence of ADHD in autumn/winter births carrying the *DRD4 7R* allele compared to the spring/summer ones with this allele. Based on those results, we hypothesized an either equal or numerically opposite pattern compared to that observed by Seeger et al. (the *winter-born ADHD hypothesis*; Figure 1b).

### Mega-analyses

We conducted *mega-analyses* – also referred to as *individual participant data (IPD) meta-analyses* – across eight studies including 3757 participants in total, to evaluate the three above-mentioned hypotheses (*the spring-focus hypothesis, the summer-born ADHD hypothesis, and the winter-born ADHD hypothesis*) in acute seasonal effects at date of measurement (*state*) and phase-sensitive learning after date of birth (*trait*), as depicted in Figure 1 (a and b). The consistency of a relationship between sleep duration and inattention between healthy individuals (4-6) and individuals with ADHD (7,8) suggests that this relationship is irrespective of diagnosis. Instead of studying the diagnosis of ADHD, we here used an approach covering inattention levels from clinical levels in ADHD patients to levels observed in healthy individuals, which is a more dynamic phenotype that allows for population level variability and may be potentially closer to the hypothesized ‘circadian pathway’ (13,14). To reduce heterogeneity, analyses focused on 18-50 years, as previous research showed a rapid shift in circadian typology from morningness to eveningness with increasing age in adolescents (36,44-47), returning back in the years after that (48). From around the age of 50, even further phase advancing is observed (48).

## MATERIALS/SUBJECTS AND METHODS

### Contributing studies

Studies were identified through the IMpACT consortium and connected research projects and gathered from August 18, 2016 until December 12, 2017. Details about the diagnostic procedures for each site (if applicable) are listed in the supplement.

The analyzed sample comprised eight studies (Table 1 and Figure 1c,1d). Each participating study had approval from its local ethics committee to perform the study and to share de-identified, anonymized individual data. Data requests for most studies required a study proposal, thereby having documented the to be included data in advance. Inclusion criteria were 1) age range between 18 and 50 years, 2) the availability of the information per participant of: a) self-rated DSM-based inattention ratings, b) *DRD4* VNTR genotyping, c) date of birth, d) the location of measurement, e) sex, f) (for the date of measurement analyses) the date of inattention rating.

**Table 1.** Overview of included studies

| Study | Country (City) | Dates of recruitment (included in current study) | Reference of main publication | Target sample | Original eligible age range (years) | Measure of inattention |
| --- | --- | --- | --- | --- | --- | --- |
| NeuroIMAGE | The Netherlands (Nijmegen, Rotterdam, Amsterdam, Groningen) | 2009-2013 | (49) | healthy participants, ADHD, unaffected siblings | 5-30 | Conners' Adult ADHD Rating Scale (CA-ARS) |
| BIG | The Netherlands (Nijmegen) | 2007-2017 | (50) | healthy participants | 19-85 | ADHD DSM-IV TR Rating Scale |
| IMpACT-Germany | Germany (Würzburg) | 2003-2017 | (51) | healthy participants, ADHD | 17-69 | Interview on DSM-IV symptoms |
| IMpACT-Brazil | Brazil (Porto Alegre) | 2002-2014 | (52) | healthy participants, ADHD | 17-68 | SNAP Inattention rating |
| IMpACT-Norway | Norway (Bergen) | 2004-2014 | (53) | healthy participants, ADHD | 19-74 | ASRS based on DSM-IV items of inattention |
| IMpACT-Spain | Spain (Barcelona) | 2002-2011 | (54) | ADHD | 16-59 | ADHD rating scale (ADHD-RS) |
| UCLA ADHD Genetics Study | United States (Los Angeles) | 1992-2009 | (55) | healthy participants, ADHD | 26-72 | ADHD rating scale (ADHD-RS) |
| MGH ADHD study | United States (Boston) | 1994-1999 | (56) | healthy participants | 18-69 | Symptom counts on the Kiddie SADs |

*IMAGE = International Multisite ADHD Genetics, BIG = Brain Imaging Genetics, IMpACT = International Multi-centre persistent ADHD CollaboraTion, UCLA = University of California, Los Angeles, MGH = Massachusetts General Hospital*

### Data selection

Individual-level data from eight studies were pooled and jointly analyzed. We applied a one-step approach, where the individual data points from all of the studies (i.e., sites) were fitted together in a single model. Each of the three hypotheses was tested in an individual model using this one-step approach. The total merged sample size was 2146 participants (1386 ADHD, 760 controls) for the test of date of measurement effects and 3757 participants (2253 ADHD, 1504 controls) for the test of date of birth effects. All participants were of Caucasian origin. Inattention ratings were z-transformed per study as well as per group (controls and ADHD patients) to harmonize different scales employed in each study and different variance distributions due to the inclusion of one or both groups per study. Sensitivity analyses were performed by fitting individual data points per study.

### Statistical Analyses

Analyses were first performed on the full sample, then split by ADHD cases and controls, and, finally, compared between these groups. The distributions of inattention measures of the total merged sample mildly violated a Gaussian distribution. Since analyses of variance (ANOVA’s) tend to be quite robust to mild violations of the Gaussian assumptions and limit false positive findings (49), we continued with these parametric tests.

Comparison of inattention between previously defined season periods – of birth or of measurement – were employed using IBM SPSS Statistics for Macintosh, version 25 (ANOVA’s). Reported effect sizes are Cohens *d*. Multiple testing was accounted for by testing three clearly defined hypotheses only. These clearly defined hypotheses allowed one-tailed testing with an alpha set on *p*=.1. Subsequent Bonferroni correction, based on the number of hypotheses, yielding a final alpha of *p*=.033. Arbitrarily, the same threshold was used for post-hoc testing further describing significant results. In addition, sexes were compared on and age was correlated with inattention to determine the need of covariation. Circannual variation was also tested using non-linear curve fitting (see supplement).

Robustness of the results was tested by comparing similarity of the circannual patterns between studies. For these tests, we used an alpha set on *p*=.1 (in this case being stricter than a lower p-value, since we here expect a *lack* of difference between studies, hence a non-significant outcome).

## RESULTS

Table 1 in the method-section describes the included studies. Table 2 provides descriptives of the participants.

**Table 2.**
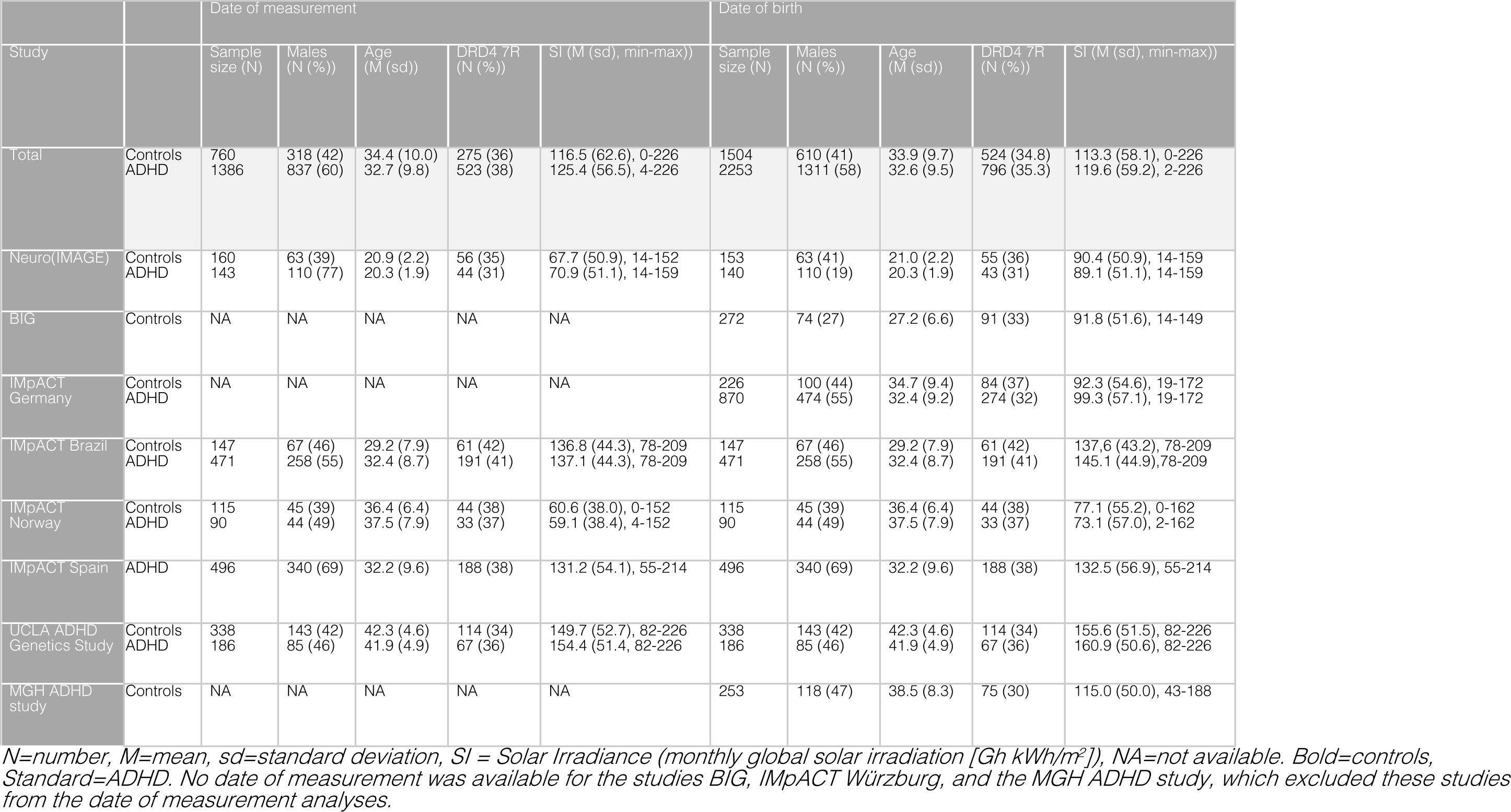
Descriptive statistics per included study.

Generally, females reported significantly more inattention (0.11±0.98) than males (−0.10±1.00) (*t*(3755)=-6.545, p<.001). Hypothesis-testing was therefore also performed with sex added as factor. Sexes did not differ on the number of 7R-carriers within the group (X^2^(1,3757)=0.681, p=.409). There was no significant correlation between age and inattention within the selected adult age-range of 18-50 years (*r*=.012, *p*=.463). Note that controlling for study (i.e. study locations) would also mean controlling for geographical differences in sunlight exposures and changes herein. Therefore, to take into account the inclusion of different sites, we conducted post-hoc sensitivity analyses.

### Spring-focus hypothesis

According to our spring-focus hypothesis (Figure 1a), we hypothesized that less inattention would be observed in 7R-carriers in the months after winter solstice, while no such change would be observed in 7R-non-carriers. We defined *“after winter solstice”* as the increasing daylength period defined by Vollmer et al. (February to April [August to October in the Southern hemisphere]) (36). *DRD4* 7R-carriers had significantly less inattention when measured *after winter solstice* (−0.173±0.970) compared to the measurements during any other time of the year (−0.017±1.023) (F(1,885)=4.744, *p*=.030, *d*=-0.156). 7R-non-carriers showed no such difference (F(1,1519)=1.526, *p*=.217, *d*=-0.071). An ANOVA between seasons (after winter solstice or not), genotype (*DRD4 7R* or non-7R), and sex (male or female) showed a significant interaction between season and genotype (F(7,2129)=4.739, *p*=.03) without interaction with sex. 7R-carriers had significantly less inattention than 7R-non-carriers after winter solstice period (F(1,722)=12.761, *p*<.001, *d*=0.271), but not in the remaining year (F(1,1420)=0.771, *p*=.380, *d*=.048). Adding diagnosis or sex to the analysis for the *after-winter solstice* period, we saw no evidence of an interaction between periods and diagnosis or sex, indicating similar outcomes for these groups.

Curve fitting results are provided in the supplement. In summary, these demonstrate a 365-day sinusoid for the period between the winter and summer solstices was preferred over a straight line particularly for 7R-carriers, for both ADHD cases and controls. Such pattern was lacking between the summer and winter solstices.

### The summer-born and winter-born ADHD hypotheses

Extrapolating from Seeger et al., (42) *DRD4* 7R-carriers born in spring/summer were hypothesized to have more inattention than those born in autumn/winter, with no such difference in 7R-non-carriers (Figure 1b). Extrapolating from Brookes et al. (43), *DRD4* 7R-carriers born in autumn/winter were hypothesized to have more inattention than those born in spring/summer, with inattention levels independent from season of birth in 7R-non-carriers (Figure 1b).

An ANOVA showed that inattention in 7R-carriers did not significantly vary between seasons (F(1,1318)=3.446, *p*=.064, *d*=-0.102). Although 7R-non-carriers had significantly less inattention among those born during spring/summer (mean=-0.015, SD=0.979) than among those born during autumn/winter (mean=0.077, SD=1.024) (F(1,2435)=5.067, *p*=.024, *d*=-0.092), this comparison lacked an a-priori hypothesis, hence require further investigation. An ANOVA between seasons (spring/summer or autumn/winter), genotype (*DRD4 7R* or non-7R), and sex (male or female) showed no significant interactions. Curve fitting results are provided in the supplement. In summary, these demonstrate no robust patterns.

### Sensitivity analyses

Because the results favor the spring-focus hypothesis, sensitivity analyses were performed to test GxE interaction effects on inattention by fitting individual data points for study-specific models (Figure 2). As can be seen in Figure 2, similar patterns were observed for the different studies. The strongest pattern was observed in the NeuroIMAGE dataset. The curve from data of this study was, therefore, compared to all of the other datasets. The relative likelihood that data from NeuroIMAGE and IMpACT Barcelona shared a curve was 87.88% (based on Akaike information criterion), statistically retaining the simple model of a shared curve (F(2,502)=0.055, p=.947), NeuroIMAGE and UCLA ADHD genetics study shared a curve with 78.66% probability (simple model retained, F(2,514)=0.726, p=.484), and NeuroIMAGE and IMpACT Brazil shared a curve with 69.58% probability (simple model retained F(2,578)=1.198, p=.303). Note that the percentage of certainty of sharing a curve decreased with the geographical location of measurement approaching the equator (where seasonal variations are less dramatically expressed). For IMpACT Norway, data points were not sufficiently well distributed throughout the year (87 7R-carriers, of which 63 with measurement day 66), which made interpretation of the observed pattern impossible.

**Figure 2.**
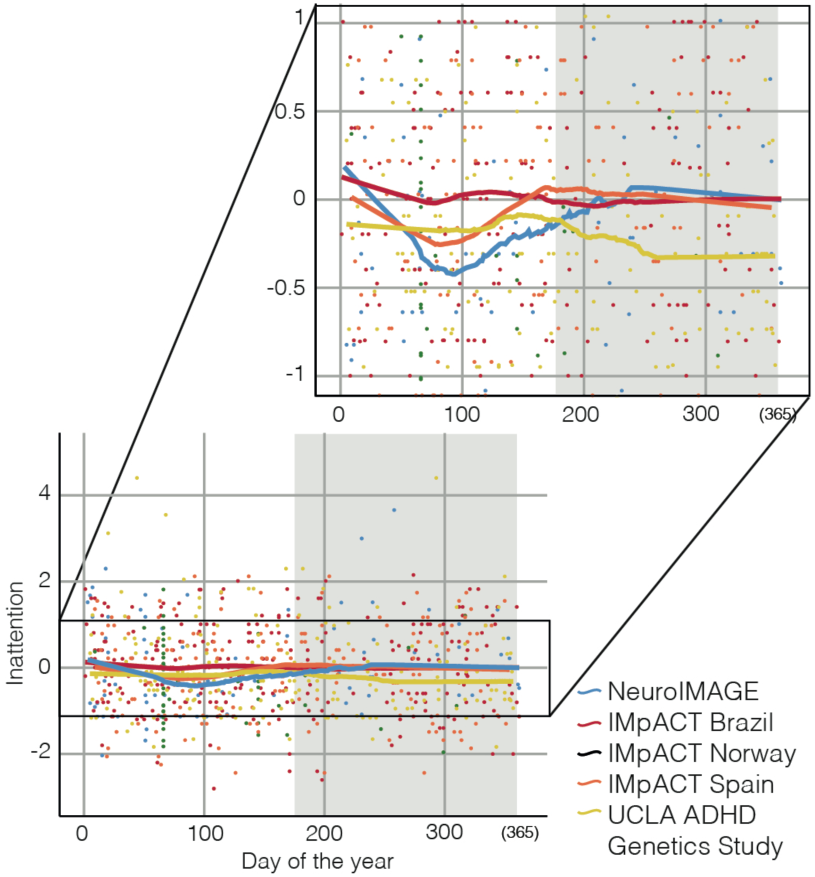
Circannual variation in inattention in 7R-carriers per study (too few data points to fit a Loess fit for Norway study). The grey shaded area indicates the period between summer- and winter solstice, the non-shaded area indicates the period between winter- and summer solstice, adjusted towards seasonality of the Northern hemisphere. A Loess Fit visualizes variation in the data by nonlinearly comparing data to its neighbouring data. Note that this procedure does not take into account the neighbouring of the last and first day of the year. NB: a more negative value implies less inattention, i.e., better attention.

#### Sunlight exposure

For each site, solar irradiation (SI) was calculated per month using “*meteonorm 7*” (http://www.meteonorm.com/en/downloads). Interpolation of data from weather stations surrounding location of the study was used (supplement, Table S1), resulting in a monthly global solar irradiation (Gh kWh/m^2^) from 7 interpolated locations used to calculate the difference between SI during the month of measurement and the preceding month [SI change (SIC)]. As expected, the standard error of the mean SIC was increasing with the amount of circannual variation observed in Figure 2 (IMpACT Brazil: SE=1.15, UCLA ADHD genetics study: SE=1.19, IMpACT Spain: SE=1.26, NeuroIMAGE: SE=1.83), illustrating larger SIC further away from the equator. When correlating SIC with the inattention ratings (Spearman-Brown due to non-Gaussian distribution of SIC), a negative correlation was found for measurements in 7R-carriers that took place during positively changing SIC (lengthening of days such as observed after winter solstice) (r=-.135, *p*=.002), whereas no such correlations were found in 7R-non-carriers or for measurements that took place during negative SIC. The larger the positive change in SI, the better the attention of 7R-carriers (Figure 3).

**Figure 3.**
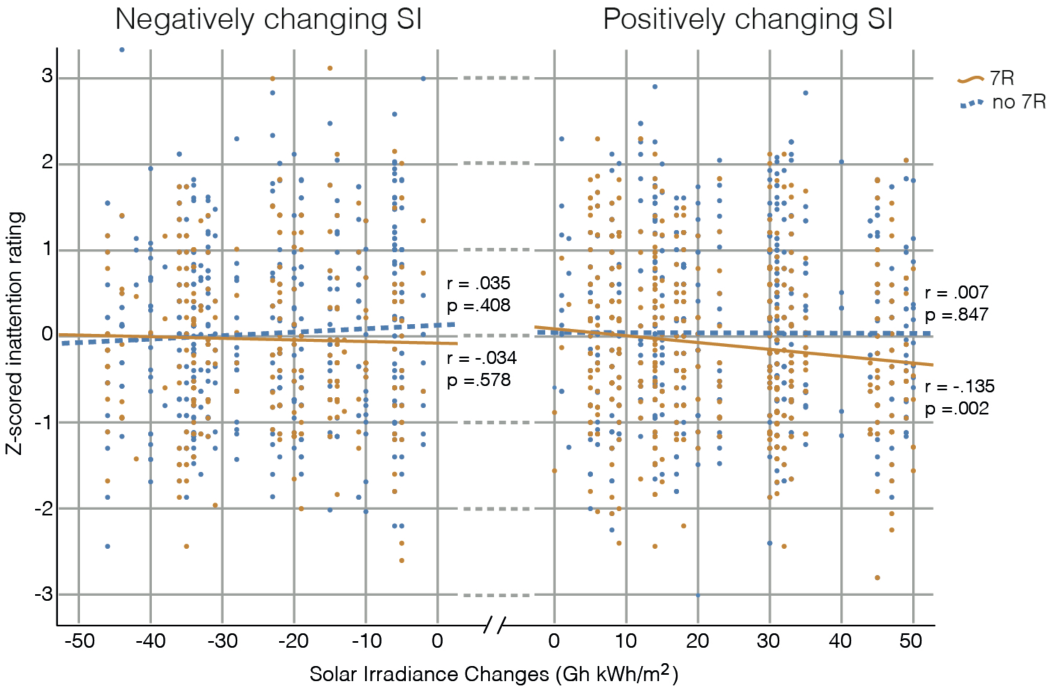
Spearman-Brown correlations between Solar Irradiance Change (SIC) and Z-scored inattention ratings for 7R-carriers (orange circles) and 7R-non-carriers (blue filled circles). NB: a more negative value implies a better outcome. All y-axes are zoomed in to [-3 – 3]. Eight data points fall outside these limits (Negative SIC, 7R-carriers: N=2, 7R-non-carriers: N=2. Positive SIC, 7R-carriers: N=2, 7R-non-carriers N=2).

## DISCUSSION

In our mega-analyses, we systematically studied inattention and the relation with date of birth or date of measurement as a function of being a *DRD4* 7R-carrier or 7R-non-carrier in participants with and without ADHD. We tested three a priori postulated hypotheses described in Figure 1. The *summer-born ADHD hypothesis*, where 7R-carriers born in spring/summer were hypothesized to have higher inattention ratings relative to those born in autumn/winter, and the *winter-born ADHD hypothesis*, where this would be opposite, were not upheld in this study. The findings of Seeger et al. (42) had already been contradicted by the findings of Brookes et al. in a much larger sample (43), and the currently observed lack of an effect is consistent with the *non-significant* findings of Brookes et al.. Our results were in accordance with the *spring-focus hypothesis*, where circadian clock response to light in 7R-carriers normalizes with increasing light exposure (difference between periods *d*=-0.156); in this case, the 7R-carriers even increased to a better level than seen in the 7R-non-carriers (difference between genotype groups *d*=0.271). The circannual pattern observed in 7R-carriers was consistent over studies, although the percentage of certainty of sharing a circannual pattern decreased with the geographical location of measurement approaching the equator (where seasonal variations are less dramatically expressed). Incorporating geographic changes into the independent measure, the change in solar irradiance at the month of measurement compared to the preceding month was determined, and was correlated to inattention in 7R-carriers. This analysis demonstrated that the larger the positive change in solar irradiance (largest in spring and furthest away from the equator), the better the attention of 7R-carriers. These results suggest that the circadian clock response to light of *less-sensitive-to-light* 7R-carriers may *sensitize* to light when its exposure increases. Albeit of note that effect sizes were small, the ability of 7R-non-carriers to reduce the light-sensitive cAMP level upon illumination (28) may be less vulnerable to such changes in environment. Effects of *DRD4* on central nervous system level cannot be ruled out at this time. We earlier also observed a lack of circannual pattern in inattention using non-genotyped data, expectedly consisting mainly of 7R-non-carriers (58). Surprisingly, 7R-carriers did not increase to a normal level after winter solstice, but to a level *better* than 7R-non-carriers. These results indicate that carrying the DRD4 7R genotype is rather beneficial for attention under certain environmental conditions. The 7R polymorphism has been proposed as one of the genetic risk factors for ADHD (22). In line with the current findings, Sánchez-Mora et al (54) had already demonstrated that the effect of the number of stressful life events on inattention scores was stronger among 7R-non-carriers than 7R-carriers. Note however, that ratings by others are needed to exclude the possibility that 7R-carriers and 7R-non-carriers report their inattention differently. Furthermore, in line with the relationship between sleep deprivation and inattention found in both healthy controls (4-6) and ADHD cases (7,8), we observed that the *DRD4 7R* genotype affected inattention regardless of ADHD diagnosis and sex.

Following the circadian pathway, we expect sleep to mediate the relationship between light exposure and inattention ratings. Still, an opposite circannual pattern to that expected based on the current pattern in inattention, was found in sleep duration in a dataset of non-genotyped healthy individuals (59). Following this pattern in sleep duration, the spring period in which we now found 7R-carriers to have better attention than in the remaining year, would be characterized by a decrease in sleep duration compared to the preceding winter period rather than an increase that would benefit attention. This discrepancy in pattern even further supports the gene specificness of the current findings.

The current findings lead to several suggestions for follow-up studies. Most relevant follow-up research would be a study where ‘the spring focus’ in 7R-carriers would be exploited. Light environment (or more specifically; changes therein) should be designed in such a way that a similar enhancement of light sensitivity is provoked as naturally occurring in February to April in the Northern hemisphere and in August to October in the Southern hemisphere. According to the current findings, such an environment should result in better attention than an environment similar to the remaining part of the year. Especially in the ADHD population, characterized by inattention that interferes with daily functioning, such exploitation would be desirable. Furthermore, the relationship between the circadian system and attentional functioning should be studied on the mechanistic level to understand its function, interaction, *and* dysfunction. Future studies of sufficient size should take into account geographically different genetic backgrounds, different versions of the circadian pathway (such as cognitive attentional performance and sleep measurements), should distinguish between adult-onset ADHD and childhood-persistent ADHD, and study other age-ranges. Unfortunately, although such data are becoming increasingly available through international efforts [e.g. Psychiatric Genomics Consortium (60)], the VNTR in *DRD4* is not often genotyped, as it is not available on genome-wide genotyping arrays. Finding (combinations of) single nucleotide polymorphisms (SNPs) that tag the 7R allele of the VNTR in genome-wide data would improve sample size and power of studies like the current one further, SNPs however do *not* mark the variation of interest (the allele length) for VNTRs when linkage disequilibrium is low in the region of the gene where the polymorphism occurs (61).

This study should be viewed in light of some strengths and limitations. Major strengths are the a priori defined inclusion criteria and generated hypotheses, preventing data mining, as well as the inclusion of the full span of inattention ratings from clinical levels in ADHD patients to levels observed in healthy populations. Also, the sensitivity analyses in which we compare data from the different studies – which can be viewed as replications – strengthens the results. Although we described a very interesting genetic susceptibility of *DRD4* 7R-carriers to seasonal changes (in daylight), the reported effects explain only a small portion of inattention ratings with small effect sizes as could be expected based on the multifactorial nature of the phenotype. We only studied inattention based on a priori hypotheses, but thereby did not study specificity of the results. For instance, large changes in solar irradiance are also associated with increased suicide attempts (62-64), and could provide an opening for further research. Furthermore, we only studied a GxE interaction, without taking into account possible gene by gene (GxG) interactions. Such interactions may be especially relevant in *DRD4* studies since dopamine *D4* and *D2* receptors are able to form heteromers and this heteromerization is influenced by genetic variants in *DRD4* (48-bp VNTR) and *DRD2* (rs2283265) (65,66). Unfortunately, very few samples have information on both DRD4 VNTR and rs2283265 precluding considering their interaction in meta-analyses. Finally, the merged sample was derived from different studies with varying study aims, methods of genotyping (albeit the different IMpACT studies and BIG all followed the same protocol), inattention operationalization, which could be viewed as both a strength (in generalizability) as well as a weakness (due to increases in variance). Conceivably, the large variance could have interfered with finding significant evidence for the hypotheses derived based on the work by Seeger and Brookes; however, it would not impugn the significant evidence for the Spring-focus hypothesis. Sensitivity analyses (Figure 2) demonstrated the consistency of the found results.

In summary, this mega-analysis demonstrated a study-consistent, diagnosis- and sex-independent, *DRD4* genotype-specific circannual variation in inattention with better ratings in periods after winter solstice (i.e. increasing solar irradiance) than other times of the year only in individuals that were *DRD4* 7R-carriers. ADHD has traditionally been described in terms of cognitive pathways (e.g., (67)). The current results, however, indicate the importance of taking into account the peripheral effects of dopamine in relation to attentional performance (mediated by the circadian clock). Direct input of midbrain dopamine to the SCN accelerates circadian entrainment, also underlining the significant relationship between the circadian system and neuromodulatory circuits related to motivational behavior (68). Better understanding of interaction between peripheral and central working mechanisms in combination with genotype-specific optimal environment conditions may provide pointers for improving treatment of inattention.

## ACKNOWLEDGEMENTS

This study made use of samples of the International Multicentre persistent ADHD Collaboration (IMpACT; www.impactadhdgenomics.com). IMpACT unites major research centres working on the genetics of ADHD persistence across the lifespan and has participants in The Netherlands, Germany, Spain, Norway, the United Kingdom, the United States, Brazil and Sweden. Principal investigators of IMpACT are: Barbara Franke (chair), Andreas Reif (co-chair), Stephen V. Faraone, Jan Haavik, Bru Cormand, J. Antoni Ramos-Quiroga, Philip Asherson, Klaus-Peter Lesch, Jonna Kuntsi, Claiton H.D. Bau, Jan K. Buitelaar, Henrik Larsson, Alysa Doyle, and Eugenio H. Grevet. We wish to thank all participants in the IMpACT studies across different countries. IMpACT is supported by grants from the departments and centers involved as well as national and international funding agencies. Among those are the European Community’s Horizon 2020 Programme (H2020/2014 – 2020) under grant agreements n° 667302 (CoCA) and 728018 (Eat2beNICE) and the ECNP Network ‘ADHD across the Lifespan’. This manuscript reflects only the authors’ views, and the European Union is not liable for any use that may be made of the information contained herein. IMpACT Norway was additionally supported by Stiftelsen Kristian Gerhard Jebsen (SKGJ-MED-002). IMpACT Spain by Instituto de Salud Carlos III (PI16/01505, PI17/ 00289, PI18/01788), and co-financed by the European Regional Development Fund (ERDF), Agència de Gestió d’Ajuts Universitaris i de Recerca-AGAUR, Generalitat de Catalunya, Spain (2014SGR1357, 2017SGR1461), the Health Research and Innovation Strategy Plan (PERIS SLT006/17/287), Generalitat de Catalunya, Spain, Departament de Salut, Generalitat de Catalunya, Spain, and a NARSAD Young Investigator Grant from the Brain & Behavior Research Foundation. Additional support was received from the European Community’s Seventh Framework Programme (FP7/2007-13) under grant agreement n° 602805 (Aggressotype)

This work also made use of the BIG database, first established in Nijmegen, The Netherlands, in 2007. This resource is now part of Cognomics (www.cognomics.nl), a joint initiative by researchers of the Donders Centre for Cognitive Neuroimaging, the Human Genetics and Cognitive Neuroscience departments of the Radboud University Medical Center, and the Max Planck Institute for Psycholinguistics in Nijmegen. The Board of the Cognomics Initiative consists of Barbara Franke, Simon Fisher, Peter Hagoort, Han Brunner, Jan Buitelaar, Hans van Bokhoven, and David Norris. The Cognomics Initiative has received support from the participating departments and centres and from external grants, i.e. the Biobanking and Biomolecular Resources Research Infrastructure (Netherlands) (BBMRI-NL), the Hersenstichting Nederland, the Netherlands Organisation for Scientific Research (NWO), and the European Community’s Seventh Framework Programme (FP7/2007– 2013) under grant agreements n° 278948 (TACTICS) and 602450 (IMAGEMEND). In addition, the work was supported by a grant for the ENIGMA Consortium (grant number U54 EB020403) from the BD2K Initiative of a cross-NIH partnership.

The NeuroIMAGE project was supported by NIH Grant R01MH62873 (to Stephen V. Faraone), NWO Large Investment Grant 1750102007010 and ZonMW Grant 60-60600-97-193 (to Jan Buitelaar), and grants from Radboud University Nijmegen Medical Center, University Medical Center Groningen and Accare, and VU University Amsterdam. M.Ribasés is a recipient of a Miguel de Servet contract from the Instituto de Salud Carlos III, Spain (CP09/00119 and CPII15/ 00023). C.Sánchez-Mora is a recipient of a Sara Borrell contract and a mobility grant from the Spanish Ministerio de Economía y Competitividad, Instituto de Salud Carlos III (CD15/00199 and MV16/00039).

Barbara Franke is supported by the Netherlands Organisation for Scientific Research (NWO) through a personal Vici grant (grant 016 130 669) and through the Dutch National Science Agenda for the NWA NeurolabNL project (grant 400 17 602).

Funding for the UCLA ADHD Genetics Study was provided by the National Institutes of Health grants MH58277and NS054124.

We wish to thank all persons who kindly participated in all the included studies.

