## Supplement for "The Role of Gene Encoding Variation of *DRD4* in the Relationship between Inattention and Seasonal Daylight"

### Procedure

*NeuroIMAGE*

The procedure followed in the NeuroIMAGE study can be found in VonRhein et al. (1). The Conners' Adult ADHD Rating Scale (CAARS) was used to rate inattention. This rating scale includes an inattention scale that is the sum of item ratings scored from 'never/rarely' [0] to 'very often' [3]. The inattention sum score was used for analyses. Genotyping was conducted analogous to Franke et al. (2).

*BIG*

Symptom counts for the ADHD symptom domains inattention and hyperactivity-impulsivity were obtained through a Dutch self-report questionnaire with 23 questions based on the DSM-IV criteria (3). Participants had to rate the occurrence (from 'never/rarely' to 'very often') of symptoms based on the past 6 months. A rating of 'often' or 'very often' was scored as a symptom. The number of scored inattention symptoms were used for analyses. Genotyping was conducted analogous to Franke et al. (2).

*IMpACT Germany, Norway, Spain*

Detailed description of the followed procedure can be found in the supplement of Franke et al. (2)

For IMpACT Germany, an interview on DSM-IV symptoms yielded the number of inattention symptoms. For IMpACT Norway, the ASRS based on DSM-IV items of inattention yielded the sum of item ratings scored from 'never/rarely' [0] to 'very often' [4]. For IMpACT Spain, the ADHD-RS was used, yielding a similar sum of item ratings, here scored from 'never/rarely' [0] to 'very often' [3]. These measures were used for analyses.

*IMpACT-Brazil*

Individuals were assessed at the adult division of the ADHD Outpatient Clinic from Hospital de Clínicas de Porto Alegre (HCPA) (cases) and at the blood bank from the same hospital (controls). All included subjects were unrelated white Brazilian adults (with 18 years of age or older) of European descent. Subjects presenting significant neurological disease that may affect cognition, head traumas, neurodegenerative disorders, history of psychosis and/or estimated IQ score lower than 70 were excluded. ADHD diagnosis was performed by experienced psychiatrists or other well-trained professionals according to the DSM-IV criteria (4) (APA, 1994) using the epidemiologic version of the Schedule for Affective disorders and Schizophrenia in semi-structured interview (K-SADS-E) (5) or the Adult ADHD Self-Rating Scale (ASRS-V1.1, (6). ADHD symptoms severity was evaluated by the Swanson, Nolan and Pelham Scale (SNAP-IV) (7). All participants signed an informed consent form approved by the Ethics Committee of the HCPA (Institutional Review Board No 0000921). The inattention rating from the SNAP-IV was used, where again a sum of inattention item ratings, scored from 'never/rarely' [0] to 'very often' [3] was calculated and used for analyses.

With respect to genotyping; DNA was extracted from peripheral blood according using salting out method, and *DRD4* polymorphism was genotyped through PCR followed by 1% synergel and 1.5% agarose gel electrophoresis.

*UCLA ADHD Genetics Study*

Participants completed the ADHD-IV Rating Scale (8) as a self-report of current ADHD symptoms, as well as an ADHD-IV Rating Scale of current ADHD symptoms in their spouse. Participants were interviewed directly using the semi-structured interview, the Schedule of Affective Disorders and Schizophrenia (SADS-LAR; (9)). The SADS-LA-IV was supplemented with the Behavioral Disorders section of the Schedule for Affective Disorders and Schizophrenia for School-Age Children – Present and Lifetime Version (K-SADS-PL) (10) to elucidate lifetime and persistent diagnoses of ADHD. The K-SADS-PL was also used to elucidate retrospective diagnosis of childhood ODD and CD. All interviews were conducted by clinical psychologists or highly trained interviewers with extensive experience in psychiatric diagnoses. 'Best estimate' diagnoses were determined after individual review of diagnoses, symptoms, and impairment level by senior clinicians. All interviews were videotaped and a subset was re-rated independently by senior clinicians to maintain ongoing reliability. The mean weighted $\kappa$ based on 24 re-rated adult tapes was 0.88 (SD = 0.10), with weighted $\kappa$ values of 0.91 for ADHD (11). The $\kappa$ values computed for initial rater against best-estimate diagnoses for 10 major psychiatric diagnoses occurring in > 5% of all subjects in the current sample was 0.95 (SD = .03).

The dopamine receptor D4 (DRD4) 48-bp VNTR was genotyped according to standard protocols (12).

*MGH ADHD study*

The families reported in this study were spouses and ADHD children of ADHD adult patients from the psychopharmacology clinic at the Massachusetts General Hospital (MGH). These adults had received a clinical diagnosis of ADHD and met DSM-IV criteria for ADHD when evaluated with the ADHD module from the Kiddie-SADS-E (13), modified to assess DSM-IV criteria. Here, the number of inattentive symptoms was used for analyses.

The spouses and children were similarly evaluated and all family members provided blood for subsequent DNA extraction. The parents provided written informed consent for themselves and their minor children who also provided written assent.

DNAs were isolated from whole blood as described by (14). The dopamine receptor D4 (DRD4) gene contains varying number of 48 bp repeats within the gene. A method described by (15) was used and modified by adding 7-deazaGTP (0.2 mM final concentration) and 10% DMSO to the PCR cocktail. The PCR products were separated by electrophoresis on 3.5% NuSieve (FMC) and stained with ethidium bromide. Laboratory procedures were blinded to the diagnoses of subjects.

### Circannual variation

Curve fitting was applied using GraphPad Prism (version 6.00 for Macintosh, GraphPad Software, La Jolla California USA, www.graphpad.com) to investigate whether circannual effects in inattention were linear or non-linear. A straight line with fixed slope and intercept of zero was compared to a sine wave with a fixed wavelength of 365 days and unconstrainted best-fits for amplitude and phase. Because a sine wave is nested in a line (i.e., a sine wave with amplitude of zero is a line), both Akaike information criteria (which uses information theory to determine how well the data supports each model) and analyses of variance

(ANOVA's; which determines how much a line model improves by chance with a sine wave model) could be derived. Circannual variation would result in a *high probability* that a sine wave is the correct model (Akaike information criterion), and *a significant deviation from chance* that a sine wave improves the model by chance (ANOVA). Curve fitting results for the main analyses are presented in the supplement.

### Sunlight exposure

For each site, solar irradiation (SI) was calculated per month using "*meteonorm 7*" (http://www.meteonorm.com/en/downloads). Interpolation of data from weather stations surrounding that site was used (Table S1), resulting in a monthly global solar irradiation (Gh kWh/m$^2$) from 7 interpolated locations used to calculate the difference between SI during the month of measurement and the preceding month [SI change (SIC)].

Table S1. *For each site (Site), the city from which Meteonorm data were derived (Interpolated city), and the exact weather stations that were included in the calculation of interpolated city (Weather stations included in 'interpolated city). Data were gathered between 1991 and 2010. The period in which the data of the nearest weather station were gathered are explicitly mentioned.*

| Study | Site | Interpolated city | Weather stations included in 'interpolated city' |
|---|---|---|---|
| **NeuroIMAGE & BIG** | **Nijmegen, Rotterdam, Amsterdam, Groningen [NL]** | Nijmegen | Wageningen (1986-2005, 21 km), De Bilt (56 km), Cabauw (BSRN) (66 km), Bochum (105 km), Zuid-Limburg/Beek (102 km), Vlissingen (162 km) |
| **IMpACT-Germany** | **Würzburg [GM]** | Wuerzburg | Wuerzburg (1996-2015, 4 km), Mannheim (106 km), Geisenheim (145 km), Weissenburg (114 km), Giessen (124 km), Stuttgart (120 km) |
| **IMpACT-Brazil** | **Porto Alegre [BR]** | - | Porto Alegre Weather Station (1991-2010) |
| **IMpACT-Norway** | **Bergen [NO]** | Bergen | Bergen/Florida (1991-2010, 0 km), BERGEN-FREDRIKSBER& (1 km), Lerwick (360 km), As (313 km) |
| **IMpACT-Spain** | **Barcelona [SP]** | Barcelona | Barcelona City (1991-2010, 5 km), Lerida/Lleida (130 km), Tortosa (154 km), Perpignan (160 km), Zaragoza Airp. (267 km) |
| **UCLA ADHD Genetics Study** | **Los Angeles [US]** | Los Angeles | Long Beach CA (1991-2005, 20 km), Los Angeles CA (22 km), SANTA BARBARA MUNI (162 km), BARSTOW-DAGGETT (157 km), San Diego Airp. CA (169 km) |
| **MGH ADHD study** | **Boston [US]** | Boston | Boston MA (1991-2005, 4 km), WORCESTER (AMOS) (67 km), Providence RI (73 km), Concord Municipal NH (103 km), Hartford/Bradley Airp (140 km) |

### Curve fitting the hypotheses

#### Spring-focus hypothesis

We tested the possible effect of *DRD4* genotypes on circannual variation observed in inattention ratings within the full sample. As can be seen in Figure 2a, a deflection of inattention ratings is seen in 7R-carriers that is lacking in 7R-non-carriers. Although this deflection is only observed in the first half of the year, a 365-day sine wave was compared to a straight line for 7R-carriers and 7R-non-carriers. In *DRD4 7R*-carriers, a sine wave was the correct model with 76.23% probability. It was significantly better than a straight line model ($F(2,796)=3.183$, $p=.042$), which shows circannual variation in inattention with date of measurement. In 7R-non-carriers on the other hand, a straight line was the correct model with 82.07% probability, retaining the null-hypothesis of a straight line ($F(2,1346)=0.486$, $p=.615$), i.e. inattention did not vary with circannual variation in this group. Hence, a model with separate curves for 7R-carriers and 7R-non-carriers was preferred with 79.98% probability and significantly improved the simple model of a shared curve for both groups ($F(2,2142)=3.393$, $p=.034$). The circannual variation in inattention with date of measurement in 7R-carriers was shared by the ADHD cases and control groups with 81.26% probability, with no statistical difference between groups ($F(2,794)=0.554$, $p=.575$).

Visual inspection of a Loess fit for 7R-carriers demonstrated particular circannual variation in the period between the winter and summer solstices. Indeed, specifically fitting data from the period between the winter and summer solstices with a 365-day sinusoid had a probability of 97.44% (rejecting a straight line, $F(2,503)=5.700$, $p=.004$, Figure S1). For the period between summer and winter solstices, a straight line was preferred with 88.05% probability (retaining a straight line, $F(2,291)=0.937$, $p=.964$). For the first period, a model with separate curves for

7R-carriers and 7R-non-carriers was preferred with 97.52% probability. This model was significantly better than the simple model of a shared curve for both groups ($F(2,1280)=5.695$, $p=.003$). For the second period, one curve for both datasets was preferred with 87.24% probability, thus, the simple model was retained ($F(2,860)=0.098$, $p=.906$). For a Loess fit through the data, see Figure 2a (Figure S1 of supplement also includes a sinusoidal fit).

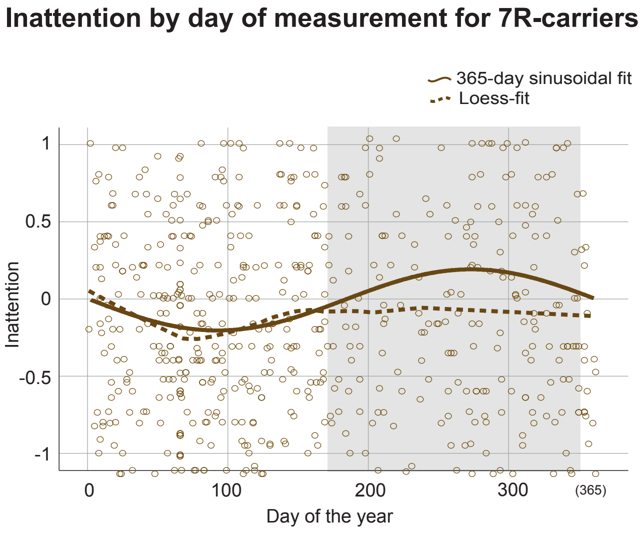


*Figure S1.* Z-scores of inattention ratings arranged by date of measurement for 7R-carriers. The grey shaded area indicates the period between summer- and winter solstice, the non-shaded area indicates the period between winter- and summer solstice, adjusted towards seasonality of the Northern hemisphere. The dashed line is a Loess-fit applied on all data, equal to that in Figure 2 of the manuscript. The solid line is a sinusoidal fit with wavelength 365, applied on the data between winter- and summer-solstice only. All y-axes are zoomed in to [-1 – 1]. NB: a more negative value implies less inattention, i.e., better attention.

### The summer-born and winter-born ADHD hypotheses

When studying inattention ratings per date of birth for *DRD4* 7R-carriers, a straight line was the favored model with 67% probability. Fitting a sine wave did not improve the model ($F(2,1318)=1.299$, $p=.273$). For 7R-non-carriers, however, results from Akaike and the ANOVA disagreed indicating non-robust results. Whereas a sine wave was the preferred model with 66.6% probability, it did not improve the straight line model ($F(2,2435)=2.695$, $p=.068$). One sine wave for both groups (ADHD cases and controls) was the favored model with 80.67% probability. Using separate models for each group did not improve the fit of the model ($F(2,3753)=0.575$, $p=.563$).

Splitting the data into participants with and without ADHD, results from the Akaike and ANOVA method disagreed again for 7R-carriers, suggesting effects were not robust. Although the model of different curves for each group (ADHD cases vs controls) was correct with 53.37% probability, this was not statistically significant ($F(2,1316)=2.146$, $p=.117$). For 7R-non-carriers, a shared curve for ADHD cases and controls was correct with 87.03% probability, and there were no significant group differences between cases and controls ($F(2,2433)=0.104$, $p=.901$). Figure 2b depicts the data with a Loess fit.

### Non-genotyped results

*Date of Measurement*

A straight line was the correct model with 84.02% probability and fitting a sine wave did not improve the model above chance level ($F(2,2144)=0.344$, $p=.709$). a shared curve for participants with and without ADHD was correct with 85.69% probability, and was retained as null-hypothesis ($F(2,2142)=0.218$, $p=.804$). Figure S2a depicts the data with a Loess fit.

*Date of Birth*

A sine wave was the correct model with 80.48% probability and significantly improved the model compared to a straight line ($F(2,3755)=3.421$, $p=.033$), demonstrating annual variation in inattention with birth dates. Most inattention was observed during the winter births. One sine wave for both groups (ADHD and controls) was the favored model with 75.20% probability. Similarly, improvement of the model when splitting the groups was at chance-level ($F(2,3753)=0.895$, $p=.409$). Figure S2b depicts the data with a Loess fit.

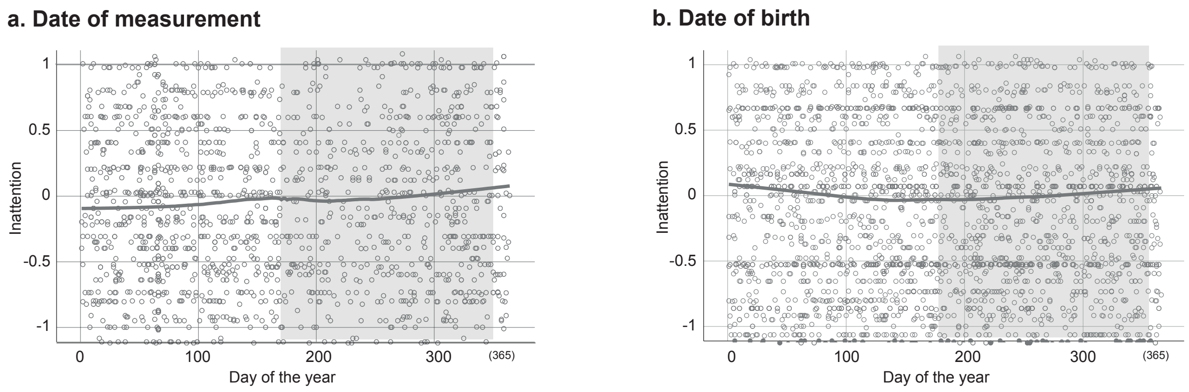


*Figure S2.* Z-scores of inattention ratings arranged by a) date of measurement or b) date of birth. The grey shaded area indicates the period between summer- and winter solstice adjusted towards seasonality of the Northern hemisphere, the non- shaded area indicates the analogous period between winter- and summer solstice. All y-axes are zoomed in to [-1 – 1]. NB: a more negative value implies less inattention, i.e., better attention.
